# Broad thermal tolerance and high mitochondrial genetic connectivity in the pinfish (*Lagodon rhomboides*)

**DOI:** 10.1101/2024.11.05.622102

**Authors:** Katherine M. Eaton, Jacob E. Samenuk, Laurel Thaxton, Victoria Chaves, Moisés A. Bernal

## Abstract

Pinfish (*Lagodon rhomboides*) are highly abundant in coastal ecosystems of the Gulf of Mexico and western Atlantic Ocean and serve as a crucial link in marine food webs. Despite their ecological relevance, little is known about this species’ susceptibility to anthropogenic climate change. Here, we characterized patterns of mitochondrial genetic divergence and examined the upper thermal tolerance of pinfish across a large portion of the species’ range. We found little evidence of population genetic differentiation among distant localities with divergent temperature regimes (e.g., Mexico and North Carolina), using two mitochondrial markers (*CytB* and *COI*). This suggests high genetic connectivity, which implies low potential for local adaptation of populations to different thermal conditions along a latitudinal gradient. To further examine population-scale differences in thermal tolerance, we assessed the critical thermal maxima (CT_max_) of pinfish from four localities: North Carolina, Florida Keys, Alabama, and Texas. We found that CT_max_ was similar across sites, with all localities showing an average CT_max_ within a 1°C temperature range (34.5-35.5°C). This suggests that southern populations of pinfish may be more susceptible to the detrimental effects of ocean warming, as individuals in lower latitudes regularly experience temperatures within a few degrees of their CT_max_. Finally, we examined the influence of varying salinity on the upper thermal limit of the pinfish and found that pinfish show no variation in CT_max_ under salinity conditions ranging from hypo-to hyper-saline (15-35 ppt). These results show that pinfish can tolerate a wide range of environmental parameters, but may rely on phenotypic plasticity rather than local adaptation to distinct conditions to cope with different environmental regimes.

## Introduction

Environmental stressors, such as changes in temperature and salinity, can directly impact physiological processes in marine organisms. Since the industrial revolution, such changes have been exacerbated by elevated concentrations of atmospheric CO_2_, which has resulted in increases in global sea surface temperature and changes in salinity regimes (IPCC, 2023; Lee et al., 2022). These fluctuations have the capacity to push marine organisms beyond their physiological limits, yet their impacts will not affect all species evenly (Hoegh-Guldberg and Bruno, 2010; Poloczanska et al., 2013; Poloczanska et al., 2016). Species’ susceptibility to climate change can be impacted by factors such as their geographic range, ecosystem stability, trophic interactions, dispersal capabilities, acclimation capacity, and adaptation potential (Poloczanska et al., 2016; Williams et al., 2008). In order to predict the long-term biological effects of climate change and implement targeted conservation strategies, it is important to begin by studying stressors at the species-level, particularly for taxa that play a disproportionate role in marine ecosystems.

The pinfish (*Lagodon rhomboides*) is a highly abundant species of forage fish found in coastal ecosystems of the Gulf of Mexico (GOM) and western Atlantic Ocean (Figure 1), ranging from Cape Cod, Massachusetts, USA to the Yucatán Peninsula, Mexico. Several aspects of this species’ biology make it an important contributor to ecological and chemical processes in coastal marine ecosystems. Larval and juvenile pinfish develop in shallow, inshore seagrass beds, and then move to deeper offshore waters as they transition to adulthood (Darcy, 1985). This movement allows for the export of nutrients from inshore estuaries and seagrass meadows to nutrient-poor offshore systems, as individuals are preyed upon by predatory fishes, seabirds, and marine mammals (Darcy, 1985; Nelson et al., 2013). The nutritional link between nearshore and offshore food webs provided by the pinfish is substantial, and it provides a major contribution to the maintenance of high-biomass offshore ecosystems of the western Atlantic (Nelson et al., 2013).

**Figure 1.**
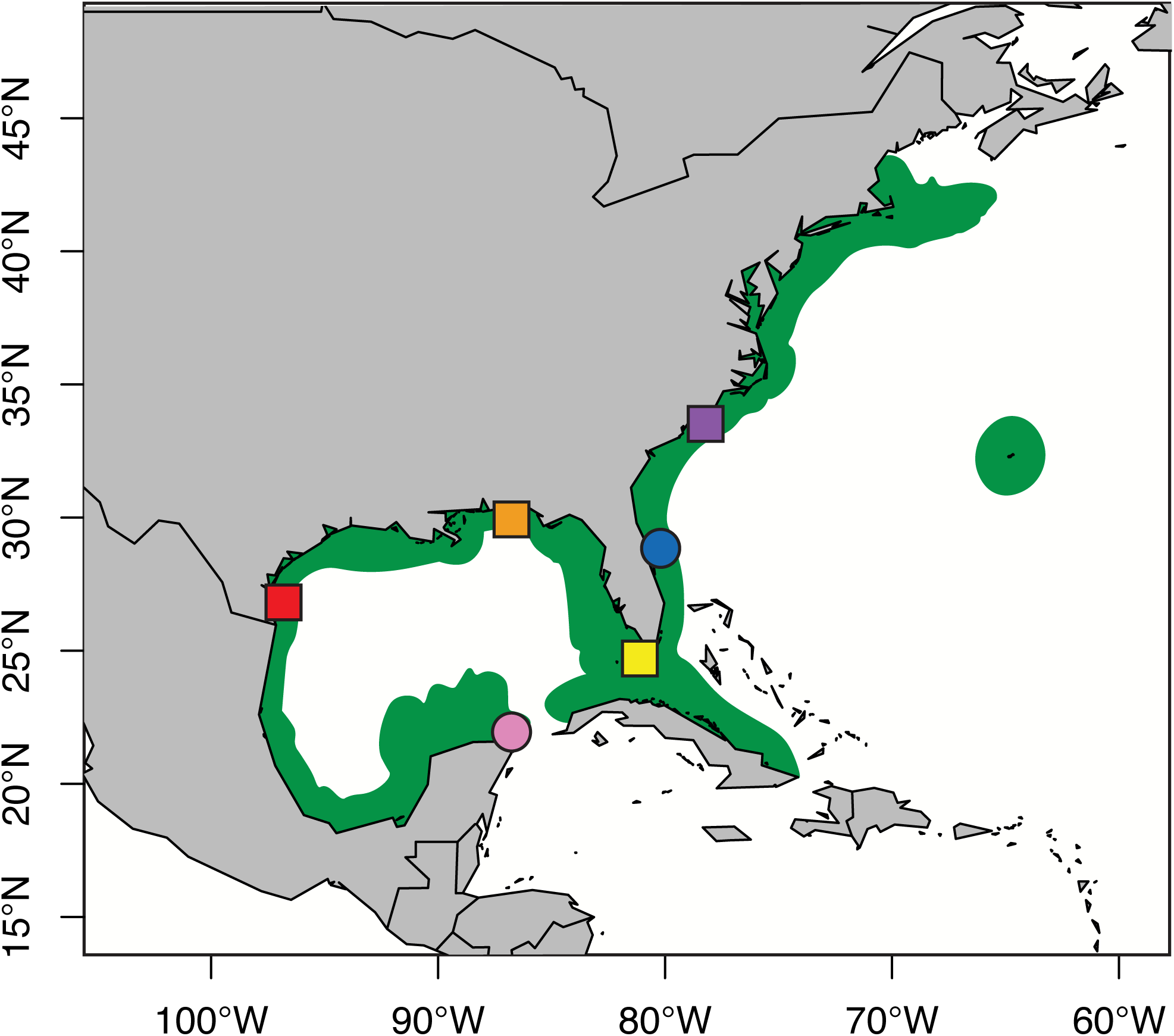
Geographic range of the pinfish (green), with sampling locations shown as colored shapes. Circles indicate locations with genetic data only (pink – Mexico; blue – Indian River Lagoon, Florida), whereas squares indicate locations with both genetic data and CT_max_ measurements (red – Texas; orange – Alabama; yellow – Florida Keys; purple – North Carolina).

Although pinfish play a crucial role in coastal Atlantic ecosystems, relatively few studies have examined the potential impacts of climate change on this species. Notably, there is a lack of published research concerning basic information on their population connectivity, as well as their thermal and salinity tolerance, which blurs our understanding of how this key species will respond to predicted changes. Previous work by Eaton et al. (2022) found that both juvenile and adult pinfish undergo increases in physiological demands and changes in gene expression in response to acute warming. However, this study examined individuals from a single population in the Northern GOM (Dauphin Island, Alabama), and did not evaluate maximum thermal tolerance (Eaton et al., 2022). Regarding salinity tolerance, Shervette et al. (2007) investigated the effects of varying salinities (0-60 ppt) on the growth and survival of juvenile pinfish from Corpus Christi, Texas, and found that salinities ranging from 15-30 ppt produced higher rates of growth and survival than extreme hypo-0 ppt) or hyper-(60 ppt) saline environments. Changes in both temperature and salinity are becoming increasingly common in the Atlantic and GOM, yet questions remain on the combined effects of both stressors on the performance and survival of the pinfish.

Maximum thermal tolerance can be heavily influenced by the latitude of a specific population, due to the potential for local adaptation to different climatic conditions (Kelly et al., 2011; Villeneuve et al., 2021). Several studies of intraspecific variation in thermal tolerance have found population-specific differences in critical thermal maxima (CT_max_), in species such as Atlantic killifish (Fangue et al., 2006), sockeye salmon (Chen et al., 2013), and redband trout (Chen et al., 2018). However, intraspecific variation in upper thermal tolerance, and the genetic connectivity between populations, remain to be comprehensively evaluated in the pinfish.

This study aims to evaluate aspects of the basic biology of the pinfish, particularly in the context of environmental tolerance and genetic divergence. Specifically, we 1) examined genetic diversity and connectivity across six populations spanning much of the pinfish’s range, 2) investigated how critical thermal maximum (CT_max_) varies by population of origin, and 3) evaluated if CT_max_ is influenced by different salinities. By assessing patterns of population differentiation and geographic variation in upper thermal tolerance, we aim to better understand the factors influencing environmental tolerance in this taxon. Ultimately, this study enhances our understanding of both genetic and phenotypic variation across the range of the pinfish and provides insights into the potential changes that coastal communities will experience in response to climate change.

## Methods

### Sample collection

Specimens of L. rhomboides from localities across the GOM and Atlantic coast were collected for analyses of population genetic differentiation. Pinfish for genetic analyses were collected via seine net from Perdido Pass and Sandy Bay, Alabama, USA, in 2020, and pinfish for salinity and thermal tolerance assays were collected via seine net from Walker Island, Alabama, USA in spring 2021 and 2023 (Alabama Department of Conservation and Natural Resources, Marine Resources Division permit numbers: 2021-3-01 and 2022-11-02). Live specimens from other localities were purchased from bait shops or procured from local fishers, after confirming the approximate collection location: n = 20 pinfish from Marathon, Florida, USA (Angler Eddy’s Live Bait & Tackle, Tavernier, FL; February 2023); n = 30 pinfish from Wilmington, North Carolina, USA (procured from local fishers in July and September 2023); and n = 31 pinfish from Corpus Christi, Texas, USA (Clem’s Marina, Corpus Christi, TX; August 2023). All subsequent experiments were done following the guidelines and under the permission of the Institutional Animal Care and Use Committee (IACUC) of Auburn University (Permits 2020-3708 and 2023-5231).

### Population genetics

DNA was extracted for 69 pinfish sampled from Texas (n = 8), Alabama (n = 36), the Florida Keys (n = 13), and North Carolina (n = 12) using the ZymoQuick DNA MiniPrep Plus Kit following the manufacturer’s instructions. Samples with low yields were re-extracted using the Invitrogen PureLink Genomic DNA Mini Kit, also per the manufacturer’s instructions.

We PCR amplified a partial sequence of the mitochondrial genes cytochrome b (CytB) and cytochrome c oxidase I (COI) from each extracted sample. Specifically, COI was amplified in a 25 μL reaction containing 0.625 U Promega GoTaq DNA Polymerase, 1X Promega Colorless GoTaq Flexi Buffer, 200 μM each dNTPs, 1.0 mM MgCl_2_, 0.4 μM each forward and reverse primer (Rocha et al., 2005; Ward et al., 2005; Table S1), and ∼100 ng DNA, and amplification conditions were: 2 minutes initial denaturation at 95°C, 40 cycles of 95°C for 30 seconds, 50°C for 45 seconds, and 72°C for 1 minute, and a final extension at 72°C for 10 minutes. CytB was amplified in a 25 μL reaction containing 0.625 U Promega GoTaq DNA Polymerase, 1X Promega Colorless GoTaq Flexi Buffer, 200 μM each dNTPs, 2.0 mM MgCl_2_, 0.4 μM each forward and reverse primer (Table S1), and ∼100 ng DNA. Amplification conditions were: 2 minutes initial denaturation at 95°C, followed by 40 cycles of 95°C for 30 seconds, 53°C for 45 seconds, and 72°C for 1 minute, and one final extension at 72°C for 10 minutes. PCR products were visually assessed via gel electrophoresis on a 1% agarose gel and sent for purification and Sanger sequencing at Azenta Life Sciences (South Plainfield, NJ).

Sequencing data was visualized in Geneious Prime v. 2023.0.4, and low-quality ends of reads were manually trimmed. Publicly available sequences for L. rhomboides were downloaded from NCBI’s GenBank and added to the analysis, to include haplotypes from Mexico (for COI; Valdez-Moreno et al., 2010) and the Indian River Lagoon (for COI and CytB; Seyoum et al., 2020; Weigt et al., 2012), and to increase our sampling depth for our sampled sites (Texas, Alabama, and the Northern Atlantic coast, for COI; April et al., 2011; Stoeckle et al., 2017; Tables S2-3). Sequences for each gene were aligned separately in Geneious Prime, using Clustal Omega v. 1.2.2. Alignments were then exported, and the IQ-Tree web server with ModelFinder was used to identify the most likely substitution model and build a maximum likelihood tree with 1,000 SH-aLRT branch test replicates and 1,000 ultrafast bootstrap replicates for each locus (Hoang et al., 2018; Kalyaanamoorthy et al., 2017; Trifinopoulos et al., 2016). Additionally, we inferred minimum-spanning haplotype networks in PopArt v. 1.7, and estimated population differentiation using Arlequin v. 3.5.2.2. Specifically, we calculated nucleotide diversity (π) and haplotype diversity (h) within and across all populations (Nei, 1987). We also used Fu’s F_S_ (Fu, 1997) and Tajima’s D (Tajima, 1989) to test for selective neutrality and population demographic history, and we calculated pairwise genetic differentiation (ϕ_st_) among populations, each with 99,999 permutations to assess significance. Finally, we conducted a hierarchical Analysis of Molecular Variance (AMOVA) to assess the relative amount of variation within populations, among populations within two major biogeographical groups (Gulf of Mexico: Mexico, Texas, and Alabama samples; Western Atlantic: Atlantic Coast, Indian River Lagoon, and Florida Keys samples), and among the two biogeographical groups, again with 99,999 permutations to assess significance.

### Thermal tolerance across populations

Live pinfish from two locations in the GOM (Texas and Alabama) and two locations on the Atlantic coast (Florida Keys and North Carolina) were transported to facilities at Auburn University and housed in 240-gallon recirculating aquarium systems, which were maintained at 21°C and 28 ppt salinity. Individuals were acclimated to life in captivity for a minimum of 14 days prior to thermal tolerance measurements.

Prior to the start of the thermal tolerance trial, fish were fasted for 12-24 hours, and then moved into the CT_max_ tanks, where they were allowed to acclimate for a minimum of 2 hours. CT_max_ trials were conducted in 8-L tanks (30 x 14.75 x 18 cm), equipped with a 1000W coil heating element (Diximus DX-1000), an Auber-WS^TM^ PID Temperature Controller (model WS-1510EBPM, Auber Instruments, Alpharetta, GA), a bubbler, and a pump for circulation. CT_max_ tanks were filled with 8 L of artificial seawater at 28 ppt. A plastic divider was used to separate the heating element, bubbler, and circulatory pump from the fish (Figure S1). Fish were tested one at a time. Trials began at room temperature (∼22°C), and water temperature was increased at a rate of 0.33°C min^-1^ until the fish displayed a loss of equilibrium (LOE). LOE is characterized by sustained uncontrolled swimming representing the point at which physiological function deteriorates and the animal loses the ability to escape from harmful environmental conditions, and it is commonly recognized as a CT_max_ endpoint (Becker and Genoway, 1979; Beitinger et al., 2000; Desforges et al., 2023). At LOE, the temperature was recorded with a precision temperature probe (Traceable Platinum High-Accuracy Thermometer, Webster, TX) with an accuracy of ±0.01℃. Once CT_max_ was reached, fish were euthanized by cervical dislocation, measured for standard length, and weighed. To evaluate the effect of location on CT_max_, a general linear model using locality and fish weight as predictor variables and CT_max_ as the response variable was performed in R v. 4.0.2. As fish weight did not have a significant effect on CT_max_ (p = 0.61), this predictor was removed from the final model, and ultimately, we performed a one-way ANOVA in R v. 4.0.2 to determine the effect of locality on CT_max_. Post-hoc corrections were applied using Tukey’s HSD.

### Effect of salinity on thermal tolerance

Live pinfish from Walker Island, Alabama were transported to facilities at Auburn University and housed in 240-gallon recirculating aquarium systems, which were maintained at 21-24℃ and 28 ppt salinity, in accordance with the seasonal average temperature and typical salinity conditions at the collection site (NOAA National Data Buoy Center, 1971). As described above, individuals were acclimated to life in captivity for a minimum of 14 days prior to CT_max_ trials.

To investigate the effects of salinity on maximum thermal tolerance, pinfish were assigned to one of five treatment groups: control (C; salinity of 28 ppt), acute low salinity (AL; salinity of 15 ppt, exposure time of 2 hours), extended low salinity (EL; salinity of 15 ppt, exposure time of 24 hours), acute high salinity (AH; salinity of 35 ppt, exposure time of 2 hours), or extended high salinity (EH; salinity of 35 ppt, exposure time of 24 hours).

CT_max_ was measured in 8-L tanks as described above, with the following modifications. For fish in the “extended” salinity treatments, approximately 24 hours prior to their CT_max_ trials, individuals were moved into a 20-gallon tank containing water of the appropriate salinity (15 ppt for the EL treatment, 35 ppt for the EH treatment). They were exposed to these salinity conditions for 24 hours, and then transferred to the 8-L CT_max_ tanks, filled with water of the same salinity (i.e., 15 ppt for the EL treatment, 35 ppt for the EH treatment), which they were then allowed to acclimate to for 2 hours prior to the start of the trial. To control for the potential effect of being moved to a novel environment (i.e., the 20-gallon tank) before being subjected to the CT_max_ trial, fish in the “control” treatment were also moved into 20-gallon tanks containing water at 28 ppt for the 24 hours prior to their CT_max_ trial (and then were likewise transferred and acclimated to water of the same salinity in the 8-L CT_max_ trial tanks). Fish in the “acute” salinity treatments were moved directly from their acclimation conditions in the 240-gallon recirculating system into the 8-L CT_max_ tanks filled with water of the appropriate salinity (i.e., 15 ppt for the AL treatment, 35 ppt for the AH treatment). Trials began at room temperature (22℃), and in this case, the temperature was increased by 0.1°C min^-1^, until loss of equilibrium was reached and CT_max_ was recorded. To evaluate the effects of salinity and acclimation time on CT_max_, a general linear model using salinity, acclimation time, and fish weight as predictor variables and CT_max_ as the response variable was performed in R v. 4.0.2. As fish weight did not have a significant effect on CT_max_ (p = 0.179), this predictor was removed from the final model, and ultimately, we performed a two-way ANOVA in R v. 4.0.2 to determine the effects of salinity, acclimation time, and the interaction between salinity and acclimation time on CT_max_.

## Results

### Population genetics

Mitochondrial gene sequencing and subsequent trimming resulted in alignments of 591 and 587 bp for COI and CytB, respectively. Genes were analyzed separately, because there were differences in the number of samples for both markers, particularly after the inclusion of publicly available sequences.

Across all populations, haplotype diversity was high for both COI (mean h = 0.8288, Table 1) and CytB (mean h = 0.9251, Table 2). Nucleotide diversity (π) was also relatively high for both genes (COI mean π = 0.004931, and CytB mean π = 0.007222, Tables 1-2). Assessing h and π values for both genes, the Indian River Lagoon population generally has the highest overall genetic diversity, whereas the Texas population generally has the lowest diversity (Tables 1-2). Tajima’s D was not significantly different from zero (p < 0.05) for any population in either gene (Tables 1-2), indicating no evidence of selection. However, Fu’s Fs was significantly less than zero (p < 0.05) for the Mexico, Alabama, and Indian River Lagoon populations, based on the COI data, and for all populations except for Texas based on the CytB data (Tables 1-2), indicating the possibility of recent population expansion based on an overabundance of private alleles.

**Table 1.** Population-wide diversity metrics for *COI* in pinfish (*Lagodon rhomboides*). Bolded values in the Tajima’s D and Fu’s Fs columns indicate values that are significantly different from the null hypothesis at p < 0.05.

| Population | Number of individuals | Number of haplotypes | Haplotype diversity (h) ± S.D. | Nucleotide diversity ($\pi$) ± S.D. | Tajima’s D | Fu’s Fs |
| --- | --- | --- | --- | --- | --- | --- |
| Mexico | 19 | 12 | 0.8947 ± 0.0589 | 0.005284 ± 0.003203 | -1.01826 | **-5.33103** |
| Texas | 9 | 4 | 0.8056 ± 0.0889 | 0.004136 ± 0.002798 | 0.46802 | 0.82623 |
| Alabama | 41 | 16 | 0.8073 ± 0.0511 | 0.004536 ± 0.002738 | -1.18003 | **-6.98992** |
| Florida Keys | 13 | 5 | 0.7308 ± 0.0963 | 0.004382 ± 0.002818 | 0.01768 | 0.57552 |
| Indian River Lagoon | 12 | 10 | 0.9697 ± 0.0443 | 0.006794 ± 0.004109 | -0.82285 | **-4.51941** |
| Northern Atlantic Coast | 17 | 7 | 0.7647 ± 0.0943 | 0.004454 ± 0.002799 | -0.04034 | -0.75132 |

**Table 2.** Population-wide diversity metrics for *CytB* in pinfish (*Lagodon rhomboides*). Bolded values in the Tajima’s D and Fu’s Fs columns indicate values that are significantly different from the null hypothesis at p < 0.05.

| Population | Number of individuals | Number of haplotypes | Haplotype diversity (h) ± S.D. | Nucleotide diversity ($\pi$) ± S.D. | Tajima's D | Fu's Fs |
| --- | --- | --- | --- | --- | --- | --- |
| Texas | 8 | 5 | 0.8571 ± 0.1083 | 0.005841 ± 0.003797 | 1.28472 | -0.00543 |
| Alabama | 36 | 23 | 0.9317 ± 0.0288 | 0.007409 ± 0.004168 | -1.32015 | **-13.81964** |
| Florida Keys | 13 | 10 | 0.9487 ± 0.0506 | 0.008103 ± 0.004761 | -0.55646 | **-3.19219** |
| Indian River Lagoon | 27 | 19 | 0.9487 ± 0.0317 | 0.008484 ± 0.004742 | -0.95372 | **-9.48470** |
| Northern Atlantic Coast | 12 | 9 | 0.9394 ± 0.0577 | 0.006272 ± 0.003839 | -0.61301 | **-3.24042** |

Pairwise estimates of population differentiation (ϕ_st_) were low across all comparisons for both markers. Notably, for COI, only two pairwise estimates of ϕ_st_ were significantly different from zero (Alabama vs. Indian River Lagoon, ϕ_st_ = 0.096 and Florida Keys vs. Indian River Lagoon, ϕ_st_ = 0.103; p < 0.05; Table 3). For CytB, none of the pairwise ϕ_st_ calculations were significantly greater than zero. However, our CytB analyses had a slightly lower sample size as there were fewer publicly available sequences for this marker (Table 4). Accordingly, the AMOVA analysis for each gene indicated that the majority of genetic variation is at the intra-population scale, rather than among populations within biogeographic regions, or among biogeographic regions (Tables S4-5).

**Table 3.** *COI*-based population pairwise differentiation (ϕ_st_) in the pinfish (*Lagodon rhomboides*), as estimated by Arlequin. Values below the diagonal indicate the ϕ_st_ value for each pairwise comparison, while values above the diagonal show the corrected p-value for that comparison. Bolded ϕ_st_ values are those which were detected as significant at p < 0.05.

| Population | Mexico | Texas | Alabama | Florida Keys | Indian River Lagoon | Northern Atlantic Coast |
| --- | --- | --- | --- | --- | --- | --- |
| Mexico | - | 0.97 | 0.95 | 0.76 | 0.07 | 0.96 |
| Texas | 0 | - | 0.87 | 0.64 | 0.16 | 0.78 |
| Alabama | 0 | 0 | - | 0.62 | 0.02 | 0.78 |
| Florida Keys | 0 | 0 | 0 | - | 0.04 | 0.51 |
| Indian River Lagoon | 0.065 | 0.042 | **0.096** | **0.103** | - | 0.06 |
| Northern Atlantic Coast | 0 | 0 | 0 | 0 | 0.075 | - |

**Table 4.** *CytB*-based population pairwise differentiation (ϕ_st_) in the pinfish (*Lagodon rhomboides*), as estimated by Arlequin. Values below the diagonal indicate the ϕ_st_ value for each pairwise comparison, while values above the diagonal show the corrected p-value for that comparison. Note that none of the pairwise ϕ_st_ estimations were significantly different from zero at p < 0.05.

| Population | Texas | Alabama | Florida Keys | Indian River Lagoon | Northern Atlantic Coast |
| --- | --- | --- | --- | --- | --- |
| Texas | - | 0.98 | 0.98 | 0.82 | 0.70 |
| Alabama | 0 | - | 0.54 | 0.37 | 0.67 |
| Florida Keys | 0 | 0 | - | 0.76 | 0.26 |
| Indian River Lagoon | 0 | 0 | 0 | - | 0.15 |
| Northern Atlantic Coast | 0 | 0 | 0.020 | 0.032 | - |

The haplotype networks for both COI and CytB revealed a relatively similar pattern of two large genetic clusters, with a smaller third cluster intermediate between the two (Figure 2). In accordance with the ϕ_st_ results, there was no genetic differentiation among populations, as each haplotype cluster contains individuals from populations across all sampled localities (Figure 2). Maximum-likelihood phylogenetic trees similarly showed no pattern of differentiation based on population of origin (Figures S2-3).

**Figure 2.**
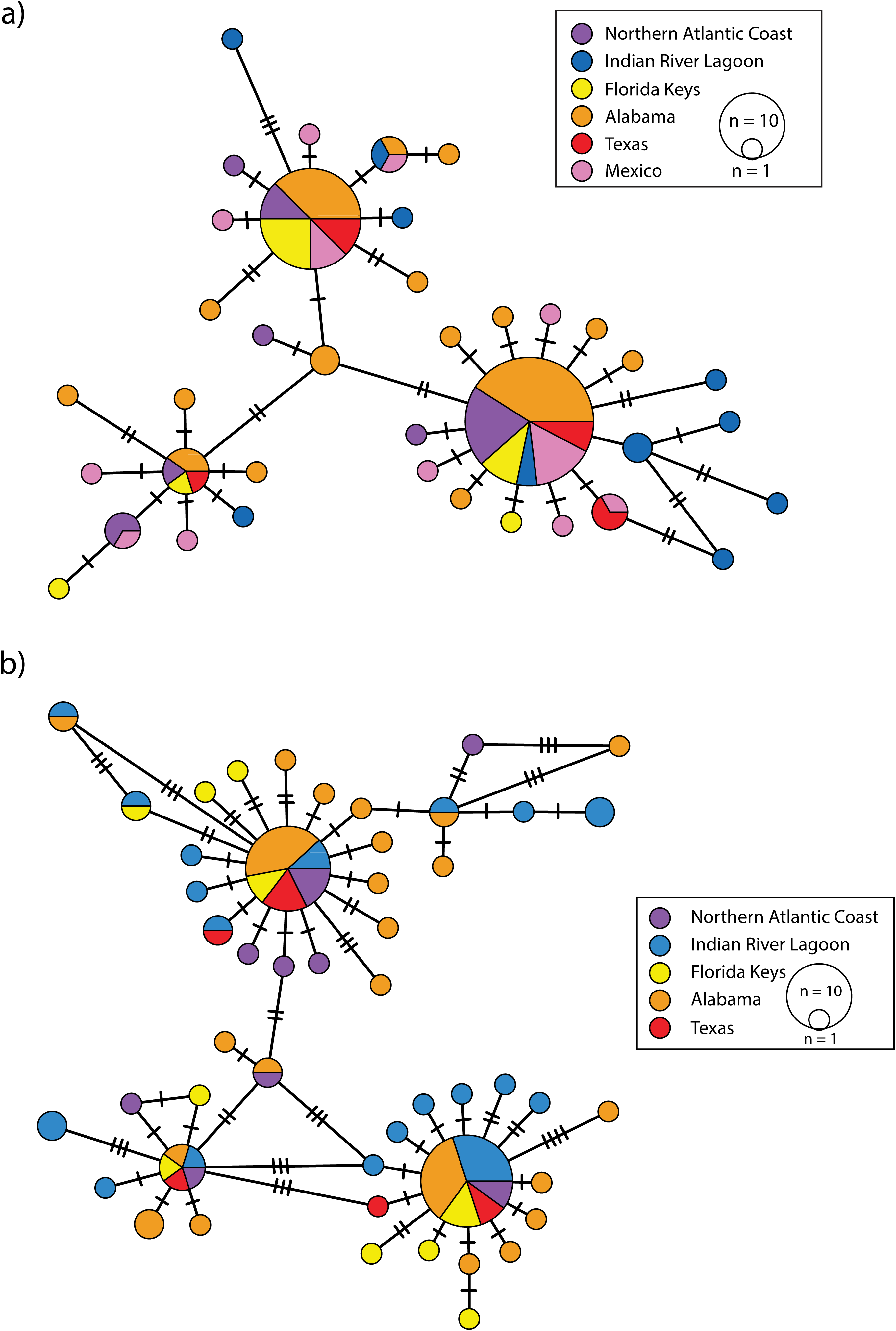
Minimum-spanning haplotype networks for the (a) *COI* and (b) *CytB* mitochondrial genes. Each circle represents a distinct haplotype, with the size of the circle corresponding to the number of individuals with that haplotype, and colored by population.

### Thermal tolerance across populations

Across populations, we found a significant effect of locality on CT_max_ (F_3,59_ = 5.722, p = 0.002), with the Alabama population having a slightly lower CT_max_ than the other three localities (Figure 3, Table S6). Specifically, Tukey’s HSD test showed that the average CT_max_ of the Alabama population was 0.67℃ lower than that of the Florida Keys population (0.11-1.24℃, 95% C.I., p-adj = 0.014), 0.70℃ lower than that of the Texas population (0.18-1.21℃, 95% C.I., p-adj = 0.004), and 0.80℃ lower than that of the North Carolina population (0.26-1.33℃, 95% C.I., p-adj = 0.001). There was no significant difference in CT_max_ among the Florida Keys, Texas, or North Carolina populations.

**Figure 3.**
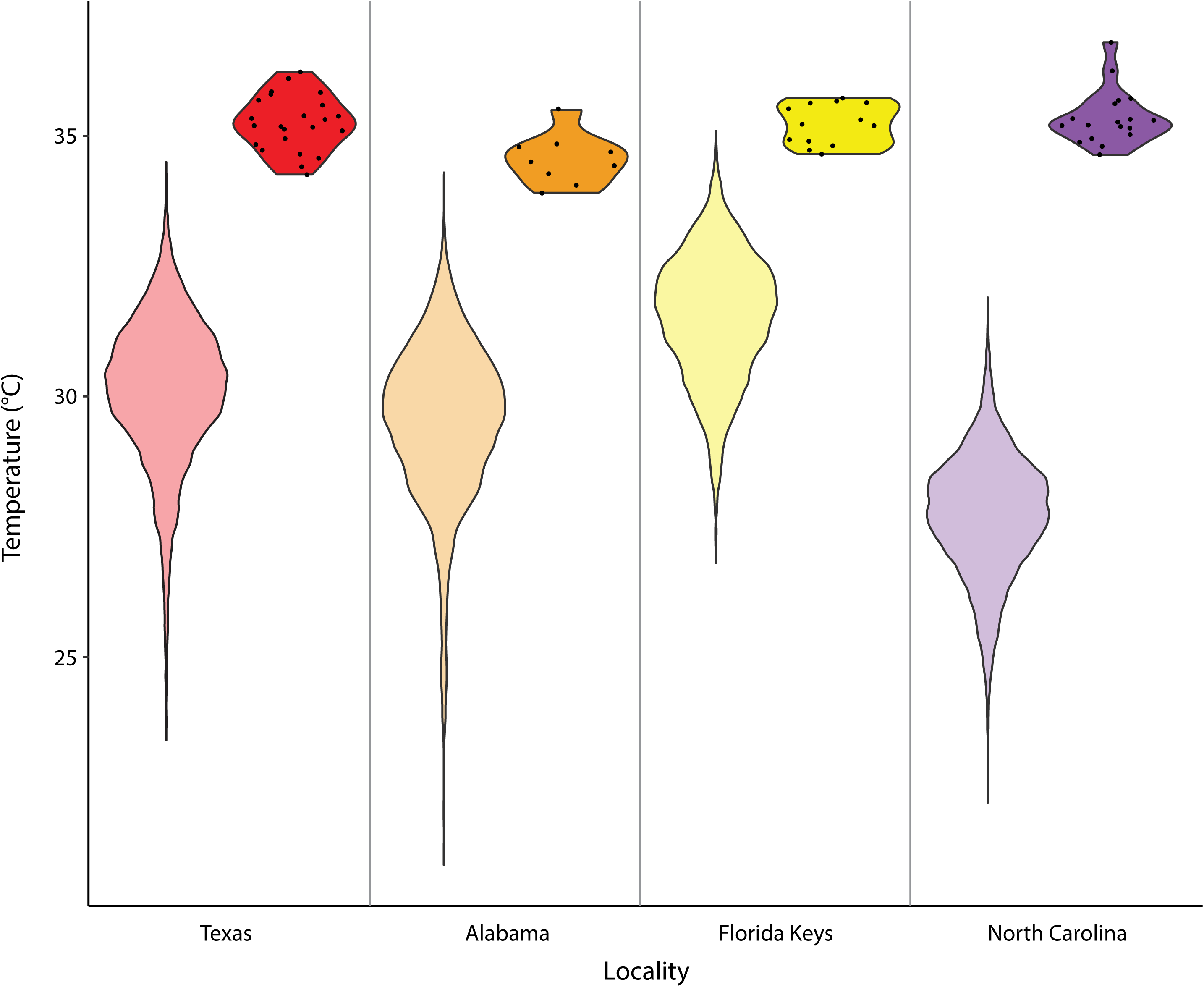
Variation in CT_max_ by locality. The lighter violin plots on the left-hand side for each locality show the distribution of recorded summer temperatures from 2014-2023 at each location (NOAA National Data Buoy Center, 1971), and the darker violin plots on the right-hand side for each locality show the distribution of measured CT_max_ values for pinfish from that locality. Individual data points for measured CT_max_ values are shown within their respective violins.

### Effect of salinity on thermal tolerance

We found that there was no significant effect of salinity (F_2,35_ = 0.452, p = 0.640), exposure time (F_1,35_ = 0.097, p = 0.757), nor the interaction between salinity and exposure time (F_1,35_ = 0.693, p = 0.411) on CT_max_ (Figure 4, Table S7). Indeed, variation in CT_max_ within treatment groups exceeded the observed variation in average CT_max_ among groups (Figure 4, Table S7), indicating that CT_max_ is not influenced by the salinity treatments induced here.

**Figure 4.**
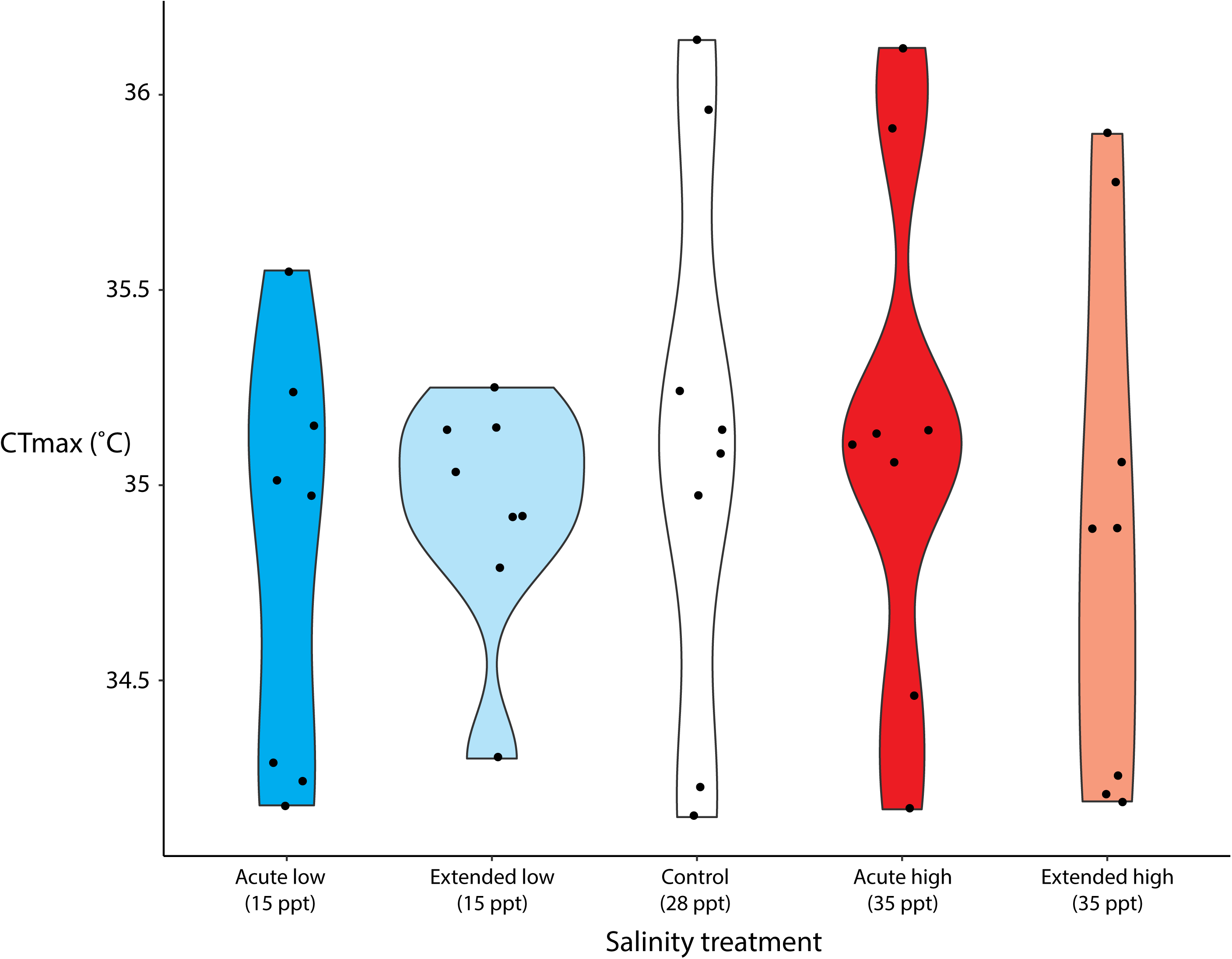
Violin plot showing variation in CT_max_ by salinity treatment. Individual data points for measured CT_max_ values are shown within their respective violins.

## Discussion

In this study, we coupled measurements of thermal tolerance with mitochondrial genetic differentiation to gain a basic understanding of the factors influencing environmental tolerance in the pinfish. We found high genetic diversity and low levels of population differentiation, and similarities in thermal tolerance across populations from different latitudes, even among distant localities with distinct climatic regimes. Additionally, we found that pinfish show no variation in upper thermal tolerance in response to changes in salinity. The widespread distribution of the pinfish, coupled with the results from this study, suggest that this species can tolerate a broad suite of environmental conditions, but that their maximum thermal tolerance has an upper limit.

### High genetic diversity and low population differentiation across the range of the pinfish

For both *COI* and *CytB*, we saw high nucleotide and haplotype diversity in each of the populations we examined (Figure 2, Tables 1-2). High levels of genetic diversity across the species’ range were coupled with low estimates of population genetic differentiation (Figure 2, Tables 3-4), as the majority of genetic variation observed was at the intra-population level (Tables S4-5). These findings indicate that there is likely high connectivity between populations, which may be due to the reproductive strategy of this species. Spawning of wild pinfish occurs offshore, and the larval stage is reported to last for at least 30 days in laboratory-reared individuals (Darcy, 1985; DiMaggio et al., 2010; Zieske, 1989). This prolonged planktonic larval duration may allow for long-distance transport via ocean currents, and ultimately a high capacity for dispersal throughout its range.

In addition to widespread population connectivity, we also found evidence supporting recent demographic expansion, as shown by the significantly negative Fu’s Fs values (Tables 1-2). The possibility of recent population growth agrees with previous estimates of demographic history based on genome-wide data, which have suggested rapid population expansion beginning around 10,000 years ago and continuing into the present day (Eaton et al., 2024).

While here, we report the first published population genetic dataset for the pinfish, future studies should focus on examining genome-wide differentiation. Mitochondrial markers such as those used in this study can be useful for reconstructing species’ phylogeography and broad-scale patterns of genetic diversity (Goodall-Copestake et al., 2012; Bowen et al., 2014). However, their lack of recombination and the maternal transmission of the mitochondrial genome place limitations on the conclusions that can be drawn regarding population differentiation and connectivity (Allendorf, 2017). Genome-wide studies of population differentiation can provide additional insight into the potential for local adaptation, even in scenarios where populations show high gene flow (Wilder et al., 2020). Furthermore, the relatively small number of individuals analyzed from certain populations here (particularly Texas, the Florida Keys, and the Indian River Lagoon) may have further impacted our results. Future studies of population differentiation in this species are needed, to provide additional insights about the potential for local adaptation in this ecologically relevant species with a broad range.

### Upper thermal limits show minimal variation among populations

In accordance with the genetic data, we saw little phenotypic differentiation among populations. Interestingly, the CT_max_ of *L. rhomboides* is similar to that of other Atlantic members of family Sparidae, such as *Sparus aurata* (35.5°C, Madeira et al., 2014) and *Diplodus sargus* (∼34°C, Madeira et al., 2012), which could indicate that common ancestry and phylogenetic relatedness plays a key role in determining a species’ upper thermal limit. Although pinfish from Alabama did have a significantly lower CT_max_ than pinfish from the other three populations, the effect size was relatively small (<1℃) and likely does not correspond to a biologically meaningful difference (Figure 3). For the most part, pinfish from different locations with distinct climatic regimes displayed relatively similar maximum thermal tolerance (Figure 3). This suggests that local adaptation of critical thermal maxima has not occurred along a latitudinal gradient in the pinfish, and that this species has evolved a broad environmental tolerance. Still, questions remain on the role that phenotypic plasticity plays for physiologically acclimating to different temperature regimes in this species.

One consideration of our findings here is that there may be an upper limit for the evolution of CT_max_ in the pinfish. The hard-ceiling hypothesis suggests that there is a firm upper limit beyond which a species cannot further increase their heat tolerance (Morgan et al., 2020). This hard ceiling could be imposed due to biochemical limitations in protein function, physiological challenges in oxygen transport and binding affinity, or changes in membrane fluidity and cellular stability beyond a certain threshold temperature (Ern et al., 2023). If this hard upper limit has been reached, this may explain the lack of differentiation in maximum thermal tolerance across sites with very different thermal regimes. It is also possible that while CT_max_ may not differ among populations, other ecologically relevant thermal performance traits may show differences. Optimal temperatures for metabolic function and reproduction may differ among populations experiencing different thermal regimes (Donelson and Munday, 2012). Further, it is possible that certain populations choose to inhabit areas within a specific thermal range, which has been considered relevant for thermal responses by ectotherms (Muñoz and Losos, 2018). The latter two considerations have not been evaluated in the pinfish, and are relevant to forming predictions about future responses to ocean warming.

Interestingly, while we found little evidence of inter-population variation in CT_max_, there is substantial climatic variation among the four localities that were sampled (Figure 3). The average summer sea surface temperature in Marathon, Florida is 31.5℃, and temperatures can regularly exceed 33℃, leaving a narrow thermal safety margin for pinfish in this locality (Figure 3; NOAA National Data Buoy Center, 1971). Conversely, in Wilmington, North Carolina, the average summer sea surface temperature is 27.7℃, and the maximum recorded temperature over the 10-year period from 2014-2023 was 31.9℃ (Figure 3; NOAA National Data Buoy Center, 1971). This suggests that pinfish at higher latitudes have much larger thermal safety margins than those at lower latitudes. As ocean temperatures continue to rise, it is possible that pinfish may undergo a northward range shift towards more suitable habitats at higher latitudes. This could have severe implications for populations in the GOM, as the North American land mass may limit the capacity of populations to expand their ranges further north (Heck et al., 2015).

### Salinity fluctuation has no effect on CT_max_

Interestingly, we found no significant difference in upper thermal limit among any of our salinity treatments (Figure 4), with most individuals having a CT_max_ around 35℃, regardless of treatment. Given that pinfish regularly occupy estuarine habitats which can undergo dramatic and rapid fluctuations in salinity, these results are not surprising. Shervette et al. (2007) reported that juvenile pinfish had high growth rates and survival when reared under salinities typical of estuarine conditions (15-30 ppt), which is approximately consistent with the conditions used in our study. This further exemplifies the broad tolerance of the pinfish to abiotic changes.

While salinity fluctuations did not affect the upper thermal limit of the pinfish measured here, it is possible that this additional abiotic stressor exacts a cost that was not detected by our phenotypic measurements. Notably, several studies have found that exposure to salinities outside of an organism’s optimum range leads to a metabolic cost in the form of increased routine oxygen demands, which increases with warming (Claireaux and Lagadère, 1999; Wuenschel et al., 2005). It is possible that a similar cost was imposed on the pinfish but was undetected by the phenotypic measurements conducted here. Future work in this species should focus on assessing changes in metabolism, growth, or survival under different combinations of salinity and temperature, to gain a more complete picture of how these two stressors may affect coastal species.

### Limitations and recommendations for future studies of upper thermal tolerance

While our assessments of CT_max_ provide a valuable resource for beginning to understand upper thermal tolerance in the pinfish, future studies should aim to address these questions at a broader scale. Our study was limited by an inability to acquire CT_max_ measurements across the entire range of the pinfish, as we were unable to include individuals in the northern-or southernmost areas of the range. As individuals at lower latitudes are most likely to be directly impacted by increasing environmental temperatures in upcoming years (Comte and Olden, 2017), it is important to assess the upper thermal limit in populations that are potentially the most vulnerable. Additionally, we were unable to assess the CT_max_ of populations north of Wilmington, North Carolina. There is a well-established biogeographic break which occurs at Cape Hatteras, North Carolina (approximately 1°N of Wilmington), due to the convergence of cooler waters from the mid-Atlantic Bight and the warm-water Gulf Stream (Briggs and Bowen, 2012; Han et al., 2022). Several studies have found phylogeographical breaks spanning Cape Hatteras in other wide-ranging species (Adams and Rosel, 2006; McCartney et al., 2013), and future work should aim to assess both genetic and phenotypic differentiation across this potential barrier to dispersal.

Further studies of upper thermal tolerance in the pinfish should also aim to implement a consistent rate of heating for determining an organism’s CT_max_. Here, because of the different heating rates used to test thermal tolerance across populations and under different salinity conditions, the results from these two experiments are not directly comparable. However, we did compare the CT_max_ results of the Alabama population (pinfish from Alabama, measured at 28 ppt salinity, with a heating rate of 0.33°C min^-1^) to those of the control salinity treatment (pinfish from Alabama, measured at 28 ppt salinity, with a heating rate of 0.1°C min^-1^). We found that, on average, fish exposed to a faster rate of heating (0.33°C min^-1^) had a CT_max_ ∼0.56℃lower than fish exposed to a slower heating rate (0.1°C min^-1^), although this result was not significant at p < 0.05 (F_1,15_ = 3.731, p = 0.07; Figure S4). It is well-established that variation in warming rate can influence an organism’s CT_max_: at very slow ramping rates, acclimation can occur, and at very rapid rates it is possible that the core body temperature of the organism may lag behind the temperature of the external environment (Beitinger et al., 2000; Desforges et al., 2023). Either of these scenarios can result in an overestimation of CT_max_ (Beitinger et al., 2000; Desforges et al. 2023). The generally recommended rate of temperature increase is 0.3°C min^-1^, and we recommend that future studies of upper thermal limit standardize to this rate. Additionally, future studies are also recommended to analyze the maximum thermal tolerance of individuals acclimated to different temperatures in aquarium settings, as acclimation temperature can also significantly impact upper thermal tolerance (Moyano et al., 2017; Strader et al., 2023).

## Conclusion

As climate change is causing major worldwide shifts in abiotic conditions, it is important to understand the factors governing environmental tolerance, particularly in ecologically relevant species. Here, we found that upper thermal tolerance in the pinfish is similar among populations across their range, with individuals from distant localities spanning a nearly 10-degree latitudinal gradient showing the same CT_max_. In accordance with this finding of little phenotypic differentiation, we found no evidence of genetic differentiation across the species’ range, which implies high genetic connectivity, potentially due to the reproductive strategy and prolonged pelagic larval phase of this species. Furthermore, we saw that pinfish appear to be robust to both acute and extended changes in salinity, showing no differences in upper thermal tolerance under hyper- or hypo-saline conditions. Ultimately, results indicate that this species is highly tolerant of an array of environmental conditions, but certain populations may be more vulnerable to climate change than others. Notably, pinfish in tropical locations such as the Florida Keys are already regularly experiencing temperatures within 1℃ of their upper thermal limit, and future warming may result in the extirpation of individuals from these localities. Future work should expand on these results using genome-wide data, which may provide finer-scale resolution and additional information regarding population connectivity and local adaptation of this important coastal species.

## Acknowledgements

We thank the members of the Bernal lab at Auburn University for helpful discussions and insight on methodology and experimental design. Kenneth Halanych, Lauren VanWoudenberg, Ally Swank, Charles Benfer, Adam Hallaj, and Diana Marchant, as well as the staff and administrators at Dauphin Island Sea Lab, the University of North Carolina at Wilmington Center for Marine Science, and the University of Texas at Austin Marine Science Institute were instrumental in assisting with sample collection. Many thanks to the Department of Biological Sciences at Auburn University and the Auburn University Undergraduate Research Fellowship Program for funding this research.

## Contributions

KME and MAB conceptualized and designed the study. KME, JES, LT, and VC carried out the experiments and analyzed the data. KME prepared the initial draft of the manuscript, and all authors contributed to the editing and finalizing of the manuscript. JES and MAB obtained funding in support of this research.

