## Supplementary Materials for "Broad thermal tolerance and high mitochondrial genetic connectivity in the pinfish (*Lagodon rhomboides*)"

**Supplemental Tables.**

**Table S1.** Sequences of primers used to PCR amplify the mitochondrial genes *CytB* and *COI*.

| Primer Name | Sequence (5’-3’) | Reference |
| --- | --- | --- |
| CytB-Forward | AATAGGAAGTATCATTCGGGTTTGATG | Rocha et al. 2005 |
| CytB-Reverse | GTGACTTGAAAAACCACCGTT | Rocha et al. 2005 |
| COI-Forward | TCAACCAACCACAAAGACATTGGCAC | Ward et al. 2005 |
| COI-Reverse | TAGACTTCTGGGTGGCCAAAGAATCA | Ward et al. 2005 |

**Table S2.** Accession numbers and localities of *COI* DNA sequences generated in this study, and publicly available sequences used to supplement population genetic analyses. Please note that while the “locality” column refers to the specific location where a sample was collected, the “population” column refers to a broader geographic region that the sample was assigned to for analyses of population differentiation.

| Accession no. | Locality | Population | Reference |
| --- | --- | --- | --- |
| GU225344; GU225635-GU225640 | Celestun, Mexico | Mexico | Valdez-Moreno et al., 2010 |
| GU224540-GU224542; GU225345-GU225348 | Quintana Roo, Mexico | Mexico | Valdez-Moreno et al., 2010 |
| JN313838-JN313842 | Quintana Roo, Mexico | Mexico | International Barcode of Life, unpublished |
| KF930020 | Brownsville, TX, USA | Texas | University of Kansas Biodiversity Institute, unpublished |
| PQ008040-PQ008047 | Corpus Christi, TX, USA | Texas | This study |
| JN026931-JN026934; HQ557326 | Sandy Bay, AL, USA | Alabama | April et al., 2011 |
| PQ008077-PQ008087 | Sandy Bay, AL, USA | Alabama | This study |
| PQ008019-PQ008026; PQ008060-PQ008076 | Perdido Pass, AL, USA | Alabama | This study |
| PQ008027-PQ008039 | Marathon, FL, USA | Florida Keys | This study |
| JQ842549-JQ842553 | Indian River Lagoon, FL, USA | Indian River Lagoon | Weigt et al., 2012 |
| MG459231-MG459237 | Indian River Lagoon, FL, USA | Indian River Lagoon | Seyoum et al., 2020 |
| PQ008048-PQ008059 | Wilmington, NC, USA | Northern Atlantic Coast | This study |
| MT455146; MT455930; MT456212 | Northampton County, VA, USA | Northern Atlantic Coast | Chesapeake Bay Barcode Initiative, unpublished |
| MT455612 | Somerset County, MD, USA | Northern Atlantic Coast | Chesapeake Bay Barcode Initiative, unpublished |
| KX688298 | New Jersey, USA | Northern Atlantic Coast | Stoeckle et al., 2017 |

**Table S3.** Accession numbers and geographic origins of *CytB* DNA sequences generated in this study, and publicly available sequences used to supplement population genetic analyses. Please note that while the “locality” column refers to the specific location where a sample was collected, the “population” column refers to a broader geographic region that the sample was assigned to for analyses of population differentiation.

| Accession no. | Locality | Population | Reference |
| --- | --- | --- | --- |
| PQ015634-PQ015641 | Corpus Christi, TX, USA | Texas | This study |
| PQ015671-PQ015681 | Sandy Bay, AL, USA | Alabama | This study |
| PQ015613-PQ015620; PQ015654-PQ015670 | Perdido Pass, AL, USA | Alabama | This study |
| PQ015621-PQ015633 | Marathon, FL, USA | Florida Keys | This study |
| MG459241-MG459250; MH352327-MH352343 | Indian River Lagoon, FL, USA | Indian River Lagoon | Seyoum et al., 2020 |
| PQ015642-PQ015653 | Wilmington, NC, USA | Northern Atlantic Coast | This study |

**Table S4.** Analysis of molecular variance (AMOVA) results for the *COI* gene.

| Source of variation | Degrees of freedom | Sum of squares | Variance components | Percentage of variation |
| --- | --- | --- | --- | --- |
| Among biogeographic regions (i.e., Gulf of Mexico vs. Western Atlantic) | 1 | 1.237 | -0.00690 | -0.48 |
| Among populations within biogeographic regions | 4 | 6.256 | 0.00812 | 0.57 |
| Within populations | 105 | 150.173 | 1.43022 | 99.91 |

**Table S5.** Analysis of molecular variance (AMOVA) results for the *CytB* gene.

| Source of variation | Degrees of freedom | Sum of squares | Variance components | Percentage of variation |
| --- | --- | --- | --- | --- |
| Among biogeographic regions (i.e., Gulf of Mexico vs. Western Atlantic) | 1 | 1.193 | -0.00913 | -0.42 |
| Among populations within biogeographic regions | 3 | 5.625 | -0.02269 | -1.04 |
| Within populations | 91 | 201.640 | 2.21583 | 101.46 |

**Table S6.** Average CT_max_ for each population (± standard deviation).

| **Population** | **Average CT_max_ (°C ± S.D.)** |
| --- | --- |
| Alabama | 34.55 ± 0.48 |
| Florida Keys | 35.23 ± 0.39 |
| North Carolina | 35.35 ± 0.52 |
| Texas | 35.24 ± 0.53 |

**Table S7.** Average CT_max_ for each salinity treatment group (± standard deviation).

| **Treatment** | **Average CT_max_ (°C ± S.D.)** |
| --- | --- |
| Control (C) | 35.11 ± 0.71 |
| Acute low salinity (AL) | 34.83 ± 0.52 |
| Extended low salinity (EL) | 34.94 ± 0.30 |
| Acute high salinity (AH) | 35.14 ± 0.65 |
| Extended high salinity (EH) | 34.90 ± 0.68 |

**Supplemental Figures.**

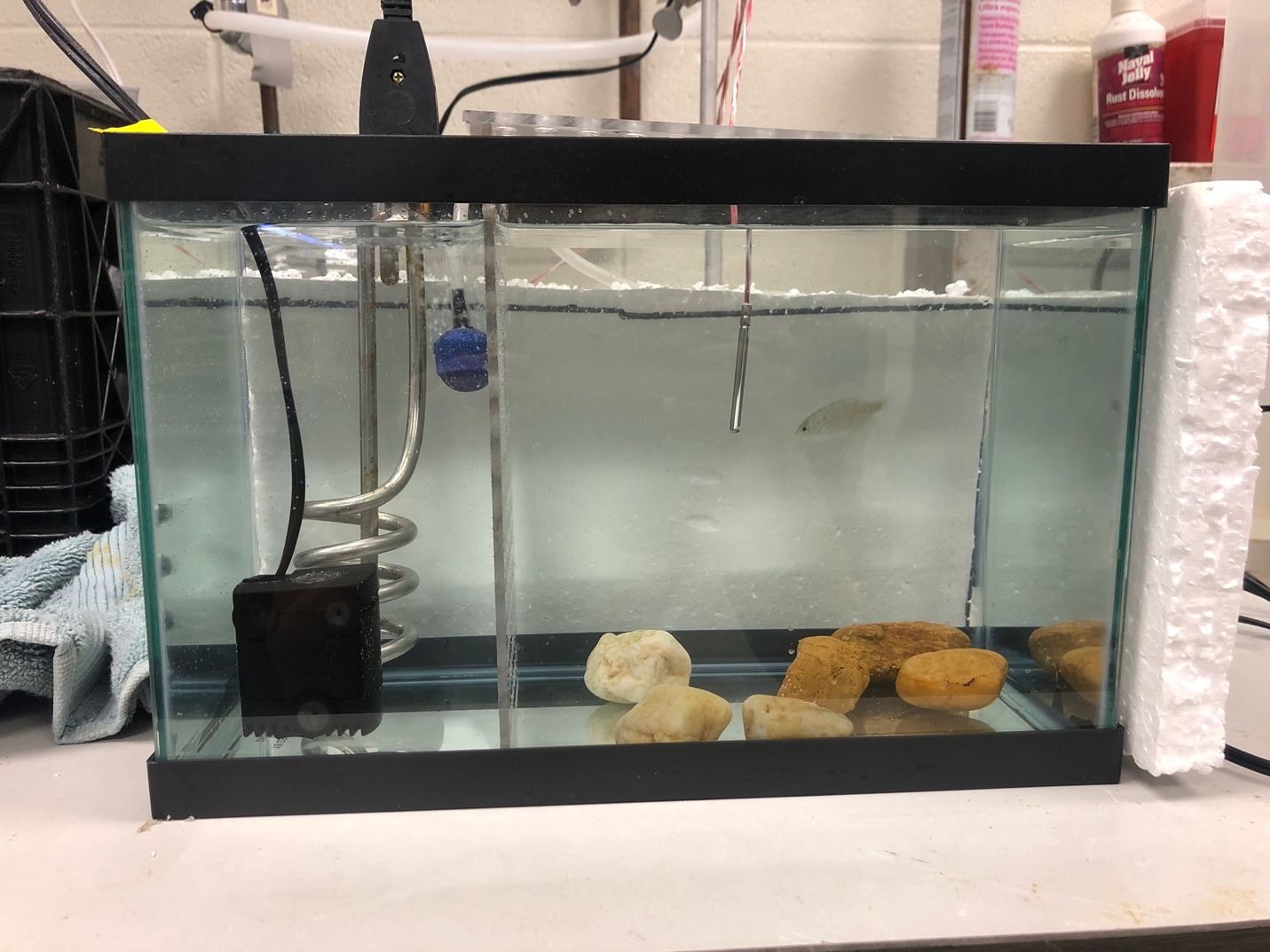

**Figure S1.** Photo of CT_max_ tank setup. The left portion of the tank houses a small pump (black), a heating element (stainless steel), and an aerator (blue). The right portion houses the fish, a temperature probe, and some rocks or PVC pipe to provide shelter and environmental complexity. The two portions of the tank are separated by a thin (~3 mm) sheet of acrylic plastic with holes drilled throughout it to allow for free flow of water between the two compartments.

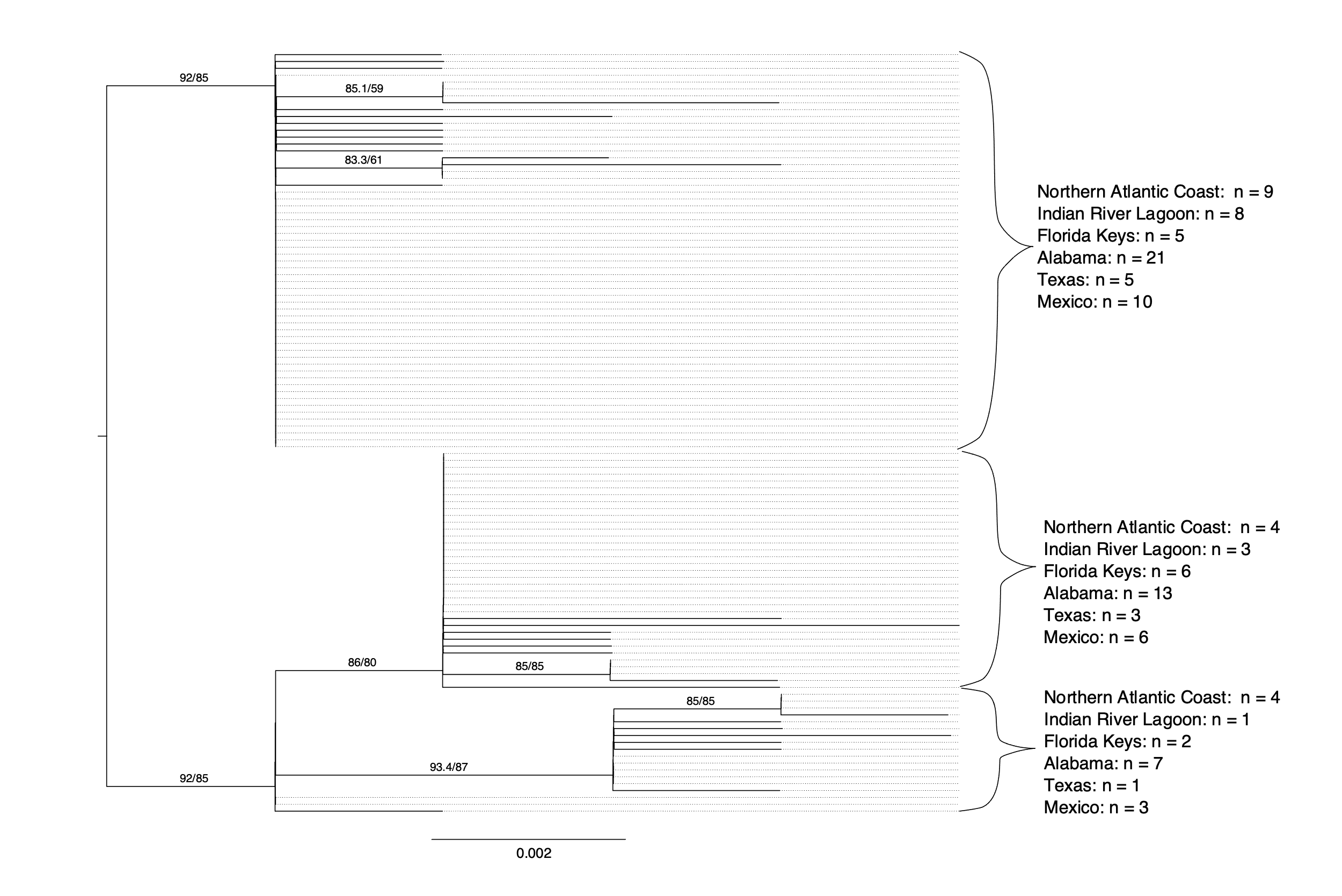

**Figure S2.** Maximum-likelihood phylogenetic tree based on sequence variation in mitochondrial gene *COI*. Support values (SH-aLRT/ultrafast bootstrap) are shown directly to the left of the node they refer to; support values lower than 80 for both tests have been removed for clarity. Branches are scaled in substitutions/site. The locality of origin for individuals in each of the three well-supported clades is shown to the right.

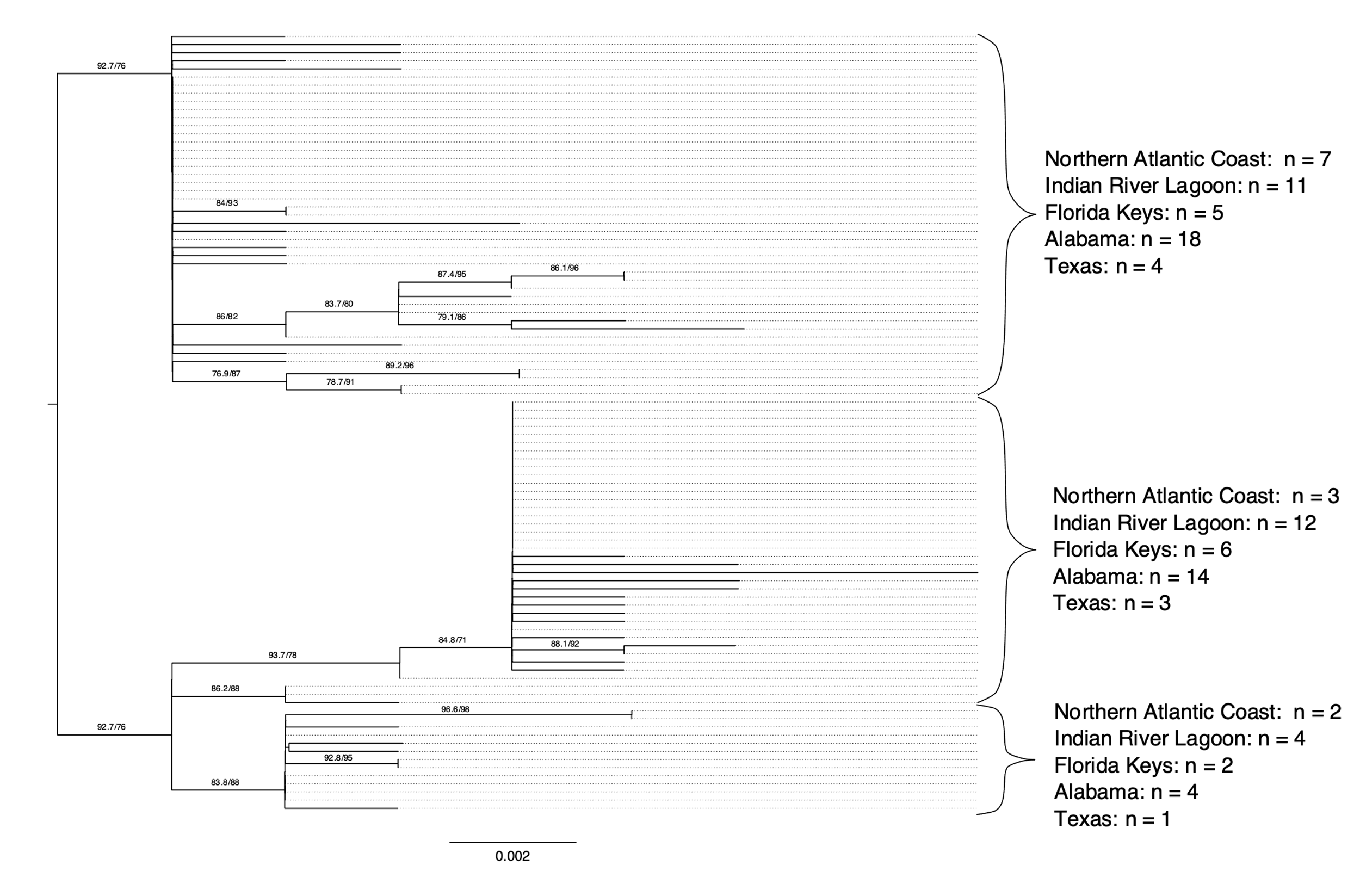

**Figure S3.** Maximum-likelihood phylogenetic tree based on sequence variation in mitochondrial gene *CytB*. Support values (SH-aLRT/ultrafast bootstrap) are shown directly to the left of the node they refer to; support values lower than 80 for both tests have been removed for clarity. Branches are scaled in substitutions/site. The locality of origin for individuals in each of the three well-supported clades is shown to the right.

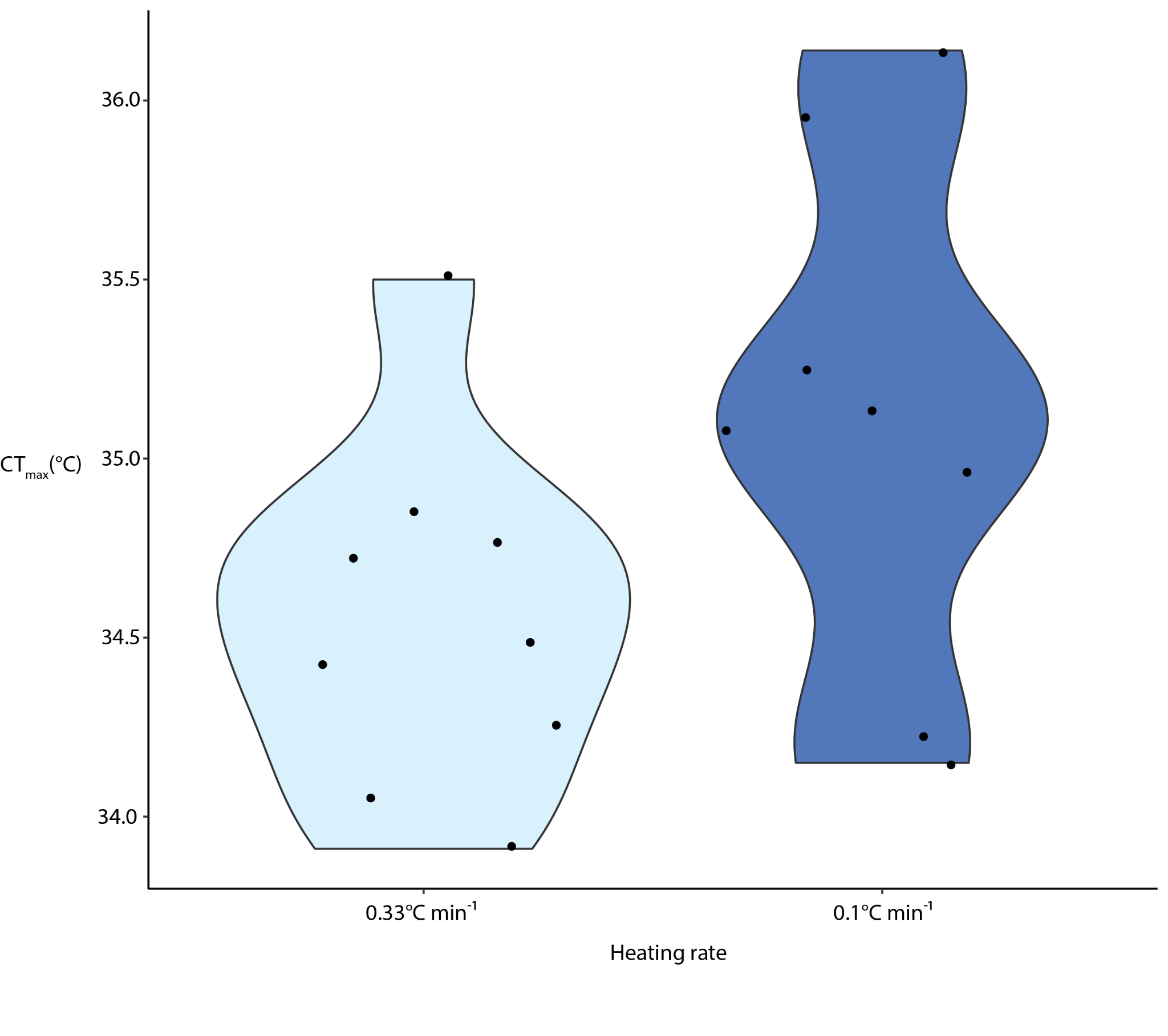

**Figure S4.** Variation in CT_max_ in individuals from Alabama when heated at the rate of 0.33°C min^-1^ vs. 0.1°C min^-1^.
